# Rearrangements and accelerated mutation rates on Dendrodorididae mitogenomes rumble their evolution

**DOI:** 10.1101/2024.06.03.597125

**Authors:** Carles Galià-Camps, Tilman Schell, Alba Enguídanos, Cinta Pegueroles, Miquel Arnedo, Manuel Ballesteros, Ángel Valdés, Carola Greve

**Author notes:** **Corresponding author**: Carles Galià-Camps. Senior researcher (Ángel Valdés, Carola Greve).

## Abstract

The systematics of the family Dendrodorididae poses a challenge to evolutionary biologists, as their mitochondrial and nuclear markers provide contradictory phylogenetic signals. Nuclear pseudogenes or exogenous contamination are hypothesized to cause the molecular discordance. However, these hypotheses have not been tested. We used genomic data from seven Dendrodorididae species to investigate the evolution of this family. Two mitogenomes displayed a novel structural rearrangement in nudibranchs, involving the translocation of three collinear genes and five surrounding tRNAs. Additionally, we found numerous mitogenomic regions with non-synonymous mutations and multiple indels in both coding and ribosomal genes. Protein modeling resulted in similar structures, suggesting that functionality is conserved. Phylogenies using mitogenomic data confirmed a specific clade membership for the rearranged mitogenomes. The incorporation of nuclear data did not fully resolve the systematic relationships of Dendrodorididae, acknowledging the evolutionary complexity of this group. The present study provides novel evidence on sudden molecular changes in mitogenomes, and highlights the relevance of using genomic data to unveil rare evolutionary processes, which is critical for understanding evolution of neglected taxa.

## INTRODUCTION

Species can be considered a set of metapopulations that evolve independently, accumulate differences, and eventually result in discrete biological units (de Queiroz, 2005). From an evolutionary perspective, the process of speciation involves several factors, with mutation and natural selection (Schluter & Rieseberg, 2022) being the main drivers. Whereas some mutations directly affect individual fitness by increasing survival under environmental pressures [3–6], others can produce phenotypes enhanced by sexual preference (Wellenreuther *et al*, 2014). Although the barriers can be reproductive or ecological, both processes eventually lead to speciation (Cooney *et al*, 2017; Mendelson & Safran, 2021; Teske *et al*, 2019; Marques *et al*, 2019). Interestingly, during the speciation process, gradients of genetic variation can be established, resulting in a range of phenotypic traits generated by recurrent gene flow between metapopulations (Abbott *et al*, 2013). Thus, speciation is mainly considered a gradual but continuous process in which a combination of processes act simultaneously to drive the divergence of metapopulations along different evolutionary trajectories, eventually leading to two new species descended from a common ancestor. (Myers, 2021; Díaz *et al*, 2018). In addition, non-adaptive mutations also have a prevalent role in speciation (Schluter & Rieseberg, 2022; Wolf *et al*, 2010; Ravinet *et al*, 2017; Xie *et al*, 2021; Wu, 2001). Drastic changes at the DNA structural level can suddenly occur, changing chromosome conformation and the interaction between their regions, such as in cases of speciation by chromosome partitioning and fission (de Vos *et al*, 2020). If the mutated individual is viable, these mutations can reduce or impede reproduction with non-mutated individuals, thus promoting accelerated speciation events through chromosome pairing incompatibilities (Řezáč *et al*, 2018).

Although widely reported, accelerated evolutionary processes driving speciation are rare (Cardini *et al*, 2007; Nowak *et al*, 2008). Genomic inversions are one of the most commonly described in the literature, in which the order of relatively long genomic sequences is reversed, hampering recombination between chromosomes (Schluter & Rieseberg, 2022; Fuller *et al*, 2019). Thus, reproductive isolation is established abruptly due to chromosome incompatibility resulting from unequal crossover (Schluter & Rieseberg, 2022; Fuller *et al*, 2019). Other mechanisms have been proposed, such as epistasis, which promotes rapid coevolution of interacting genes that can accomplish a high level of genetic differentiation in a few generations (Breen *et al*, 2012). High evolutionary rates can produce incompatibilities among individuals of the same species, eventually generating different evolutionary units (Lecturer Faculty of Life Sciences Jason B Wolf *et al*, 2000; Publication times & Open access; Phillips, 2008). The nuclear genome is not the only one responsible for fast-occurring incompatibilities since similar processes can occur in the mitogenome (Burton, 2022; Tobler *et al*, 2019; Hill, 2016). Cyto-nuclear interactions can play an important role in speciation since genes from both genomes are necessary for the correct functioning of cellular respiration (Sutovsky, 2019), as well as for an accurate and even distribution of mitochondria during mitosis and meiosis (Ghiselli *et al*, 2019). Thus, mutations occurring in one of these interacting genomic regions must be repaired or compensated by the counterpart through a counter-mutation (Hill, 2020). Once the compensation process occurs in a specific lineage, mitochondrial and nuclear mutations can cause incompatibilities with other individuals from the same species (Burton, 2022; Visinoni & Delneri, 2022).

The family Dendrodorididae, which includes the genera *Dendrodoris* and *Doriopsilla*, presents several challenges for integrative taxonomists. Members of this family lack taxonomically important structures present in most other gastropods, such as shells, radula, and jaws, and their mitochondrial and nuclear markers provide contradictory phylogenetic signals (Galià-Camps *et al*, 2022). Additionally, phylogenies reconstructed with partial cytochrome oxidase I (*cox1*) and ribosomal 16S sequences display an extremely long branch for some species of the genus *Dendrodoris* (Hallas *et al*, 2017; Korshunova *et al*, 2020). Long branches in phylogenies usually reflect distant evolutionary relationships, although they might indicate accelerated evolutionary rates when found in closely related taxa (Omilian & Taylor, 2001). Nevertheless, *Dendrodoris* long branches have been attributed to contamination or a pseudogene randomly resembling those of insect *cox1* as found with BLAST (Hallas *et al*, 2017; Korshunova *et al*, 2020). However, neither the pseudogenes nor the contamination hypotheses have ever been tested.

Here, we use low-coverage whole genome sequencing of seven Dendrodorididae species to recover their mitogenomes and generate their first nuclear genome draft. We validate the functionality of mitochondrial coding sequences (CDS) by modeling their tertiary structure. We use mitochondrial gene sequences independently and in combination with nuclear single-copy ortholog genes to assess the family phylogenetic relationships. Furthermore, we use genome-wide markers to validate the results of the phylogenetic analyses. With the results presented here, we discuss the evolutionary processes that may have shaped the mitogenome structure of Dendrodorididae and the phylogenetic relationships of the family. The present study presents novel results on drastic changes in molluscan mitogenomic evolutionary processes, discusses the cyto-nuclear interplay, and explores possible respiration adaptations produced by habitat change.

## RESULTS

### Mitogenomic and nuclear properties

Illumina whole genome sequencing produced between 8.67Gb and 40.34Gb (Table S1). After data curation, we kept from 5.96 to 38.91Gb (Table S1). We successfully circularized and annotated the mitogenomes of the seven Dendrodorididae representatives. All mitogenomes had the regular composition of 13 CDS, 2 rRNA, and 22 tRNA with high and continuous coverage (Table S1). Smooth *Dendrodoris* species had the smallest mitogenome sizes, followed by the members of the genus *Doriosilla*, and finally warted *Dendrodoris* and Phyllidididae species with similar values (Figure 2A, Table 2). For GC content, smooth *Dendrodoris* had the smallest values, followed by *Doriosilla*, Phyllidididae, and warted *Dendrodoris* (Figure 2A).

**Figure 1:**
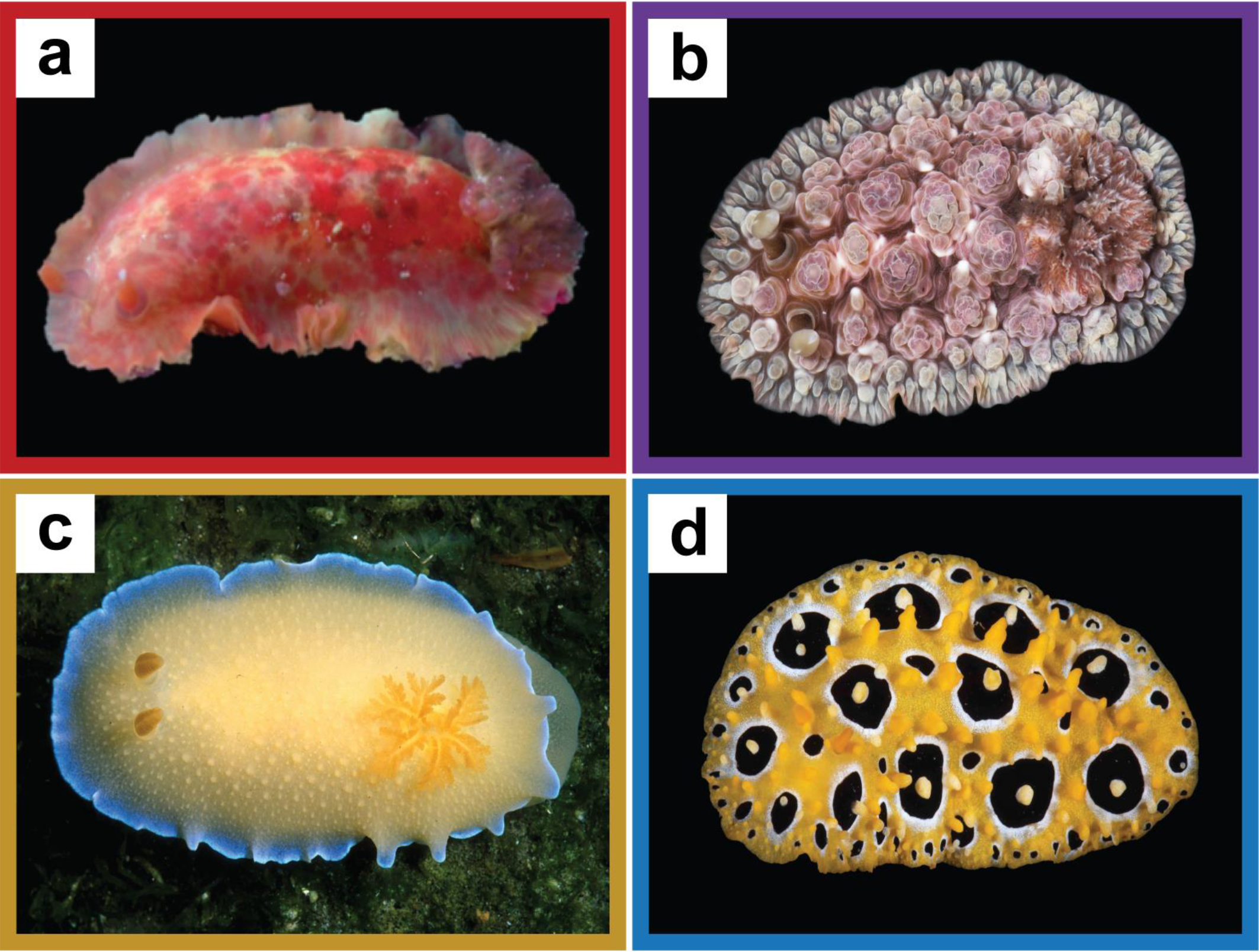
Pictures of a representative species for each taxon used in the present study. **a)** Smooth *Dendrodoris: Dendrodoris temarana*. Source: Carles Galià-Camps **b)** Warted *Dendrodoris: Dendrodoris tuberculosa*. Source: Ángel Valdés **c)** *Doriopsilla: Doriopsilla spaldingi*. Source: NHM Los Angeles County **d)** *Phyllididae: Phyllidia ocellata.* Source: Ángel Valdés

**Figure 2:**
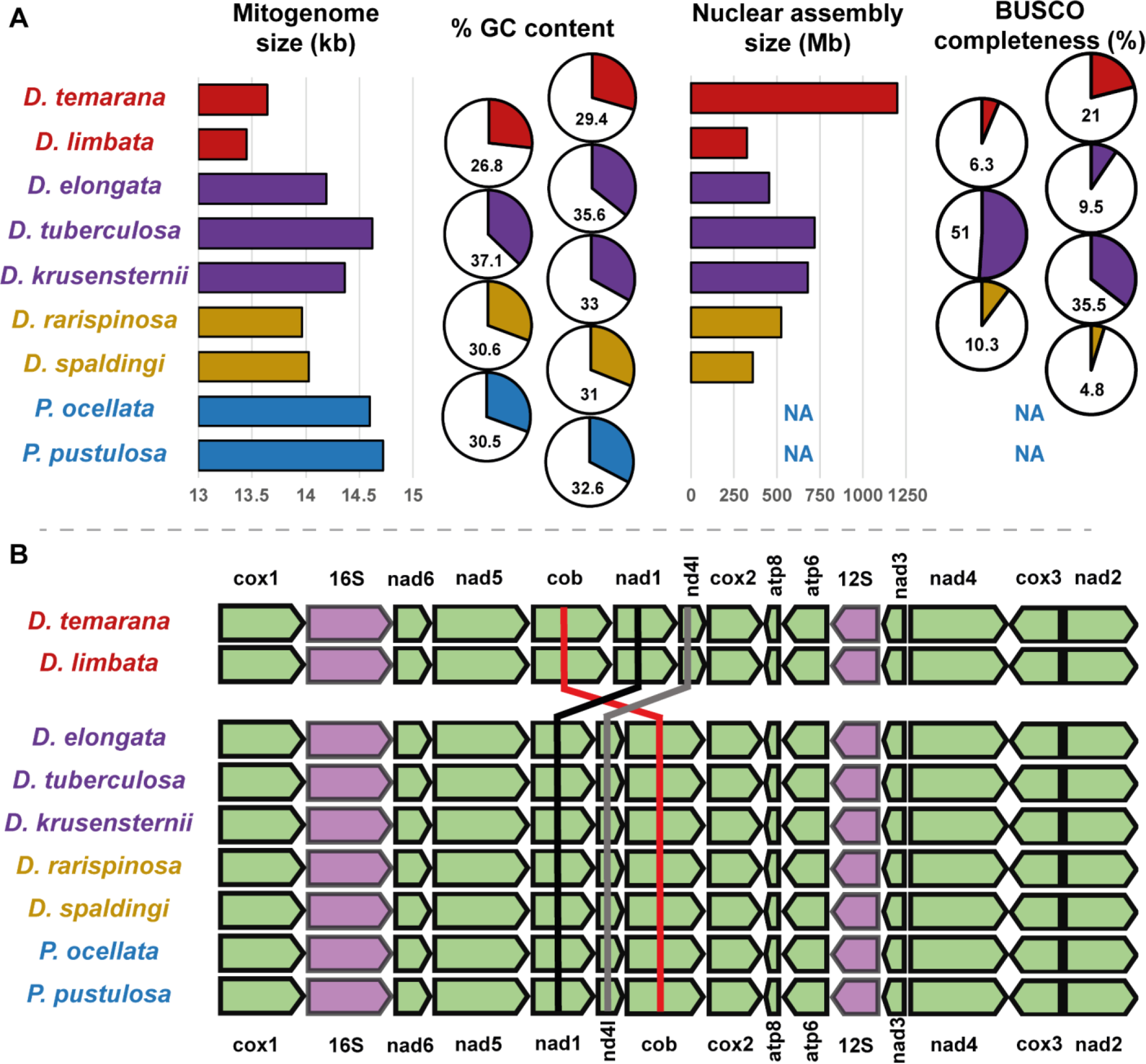
Mitogenomic features of smooth *Dendrodoris* (red), warted *Dendrodoris* (purple), *Doriopsilla* (yellow), and the outgroup species Phyllididae (blue). **A)** For each species, mitogenome assembly size (kb), %GC content, nuclear assembly size (Mb), and BUSCO completeness (%). Note that nuclear features for Phyllididae species are not available. **B)** Mitogenomic gene order for each species studied. Black, grey, and red lines are displayed for *nad1*, *nd4l,* and *cob* respectively to facilitate mitogenome rearrangement interpretation.

Nuclear assemblies ranged from 325Mb (*Dendrodoris limbata*), to 1200Mb (*D. temarana*) (Table S1). Overall, BUSCO completeness varied with genome length, ranging from 4.8% to 51%. (Table S1). Considering the genome length of 700Mb and the BUSCO completeness of 51%, we defined the genome of *D. tuberculosa* as the most comprehensive pseudo-reference genome assembly for nuclear SNP analyses.

### Mitogenomic sequences and genes

Alignment of the mitogenomes revealed a translocation involving the *cob*, *nad1* and *nad4l* genes, which was specific for the smooth *Dendrodoris* morphotype (Figure 2B). The translocation also involved the relocation of 5 surrounding tRNAs. Single gene alignments presented many mutations when comparing smooth *Dendrodoris* against all other sequences at both the nucleotide and amino acid levels. There were many insertions and deletions (indels) for the rRNA genes, most specific to the smooth *Dendrodoris* species morphotype (Figure 3A). CDS alignments of amino acids also displayed indels (Figure 3A), differentiating smooth *Dendrodoris* from the other taxa. *Dendrodoris* showed atypically high values for both synonymous and non-synonymous changes. Interestingly, d*N*, dS and d*N*/d*S* values comparisons involving smooth *Dendrodoris* (d*N*=1.529 ± 2.840, d*S*=57.228 ± 48.396, d*N*/d*S*=0.273 ± 2.496; mean ± standard deviation) were much higher than without them (d*N*=0.184 ± 0.172, d*S*=42.535 ± 35.684, d*N*/d*S*=0.009 ± 0.014) (Figure 3, Supp Spreadsheet 1). GLMs for d*N*, d*S*, or the d*N*/d*S* ratio values confirmed that gene and sequence interaction was significant (Table 1). For d*N*, smooth *Dendrodoris* comparisons were always significantly different regardless of the gene (Figure 3, Supp Spreadsheet 2). For d*S*, only *cob* and *cox1* presented significant differences between comparisons involving smooth *Dendrodoris* and without involving them. For d*N*/d*S*, *cox2*, *cox3*, *nad1*, *nad2*, *nad3*, *nad4l*, and *nad6* showed differences between comparisons (Figure 3, Supp Spreadsheet 2).

**Figure 3:**
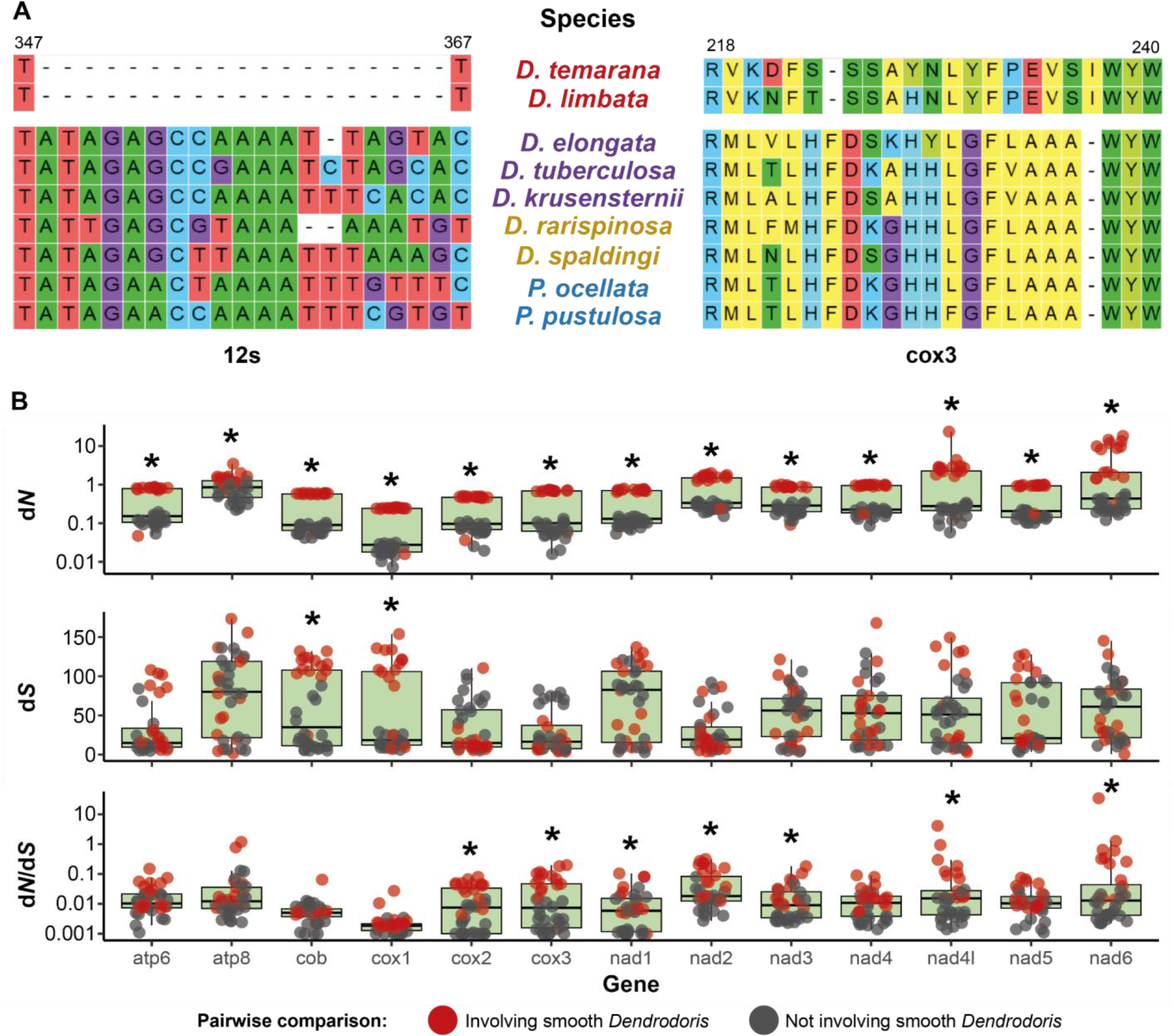
Mitochondrial genes comparisons. A) Examples of gene alignment and presence of smooth *Dendrodoris* lineage-specific indels on ribosomal genes (left panel; *12S*) and protein-coding genes (right panel; *cox3*) **B)** Sequence pairwise comparison values for d*N*, d*S*, and d*N*/d*S*. Asterisks above the boxplots indicate significant differences (p<0.05) after applying a t-test between comparisons involving and not involving smooth *Dendrodoris* sequences.

**Table 1:** GLM results for the dN, dS, and dN/dS values. Note that all values were log10 transformed. As fixed factors we included Comparison (involving smooth Dendrodoris, not involving smooth Dendrodoris) and Gene (cox1, cox2, cox3, cob, atp6, atp8, nad1, nad2, nad3, nad4, nad4l, nad5, nad6). For each factor and their interaction, we provide their sum of squares (Sum Sq), degrees of freedom (DF), F value, and p-value. Significant p-values are in bold.

| Model | Factor | Sum Sq | Df | F | p-value |
| --- | --- | --- | --- | --- | --- |
| dN | Comparison | 27.00 | 1 | 98.63 | **<0.001** |
|  | Gene | 177.37 | 12 | 54.00 | **<0.001** |
|  | Comparison*Gene | 24.97 | 12 | 7.60 | **<0.001** |
|  | Residuals | 120.98 | 442 |  |  |
| dS | Comparison | 6839.00 | 1 | 5.15 | **0.024** |
|  | Gene | 78061.00 | 12 | 4.90 | **<0.001** |
|  | Comparison*Gene | 118612.00 | 12 | 7.45 | **<0.001** |
|  | Residuals | 586778.00 | 442 |  |  |
| dN/dS | Comparison | 8.47 | 1 | 7.36 | **0.007** |
|  | Gene | 96.80 | 12 | 7.01 | **<0.001** |
|  | Comparison*Gene | 90.33 | 12 | 6.54 | **<0.001** |
|  | Residuals | 508.69 | 442 |  |  |

**Table 2:** Pairwise comparison of mitochondrial protein structure models. For each protein, species being compared are indicated in columns Taxon 1 and Taxon 2. For each comparison, we provide sequence identity (%), root-mean-square deviation of atomic positions in armstrongs (RMSD Å), and template modeling score (TM-score).

| Protein | Taxon 1 | Taxon 2 | Sequence identity (%) | RMSD (Å) | TM-score |
| --- | --- | --- | --- | --- | --- |
| atp6 | smooth *Dendrodoris* | warted *Dendrodoris* | 28 | 1.12 | 0.94 |
|  | smooth *Dendrodoris* | *Doriopsilla* | 30 | 1.14 | 0.94 |
|  | warted *Dendrodoris* | *Doriopsilla* | 68 | 1.55 | 0.94 |
| atp8 | smooth *Dendrodoris* | warted *Dendrodoris* | 19 | 1.57 | 0.37 |
|  | smooth *Dendrodoris* | *Doriopsilla* | 13 | 2.23 | 0.06 |
|  | warted *Dendrodoris* | *Doriopsilla* | 0 | 1.8 | 0.12 |
| cob | smooth *Dendrodoris* | warted *Dendrodoris* | 42 | 0.88 | 0.94 |
|  | smooth *Dendrodoris* | *Doriopsilla* | 44 | 0.87 | 0.96 |
|  | warted *Dendrodoris* | *Doriopsilla* | 85 | 0.61 | 0.97 |
| cox1 | smooth *Dendrodoris* | warted *Dendrodoris* | 66 | 0.33 | 0.99 |
|  | smooth *Dendrodoris* | *Doriopsilla* | 65 | 0.3 | 0.99 |
|  | warted *Dendrodoris* | *Doriopsilla* | 95 | 0.14 | 0.99 |
| cox2 | smooth *Dendrodoris* | warted *Dendrodoris* | 48 | 0.07 | 0.99 |
|  | smooth *Dendrodoris* | *Doriopsilla* | 46 | 0.07 | 0.99 |
|  | warted *Dendrodoris* | *Doriopsilla* | 85 | 0.04 | 1 |
| cox3 | smooth *Dendrodoris* | warted *Dendrodoris* | 41 | 0.11 | 1 |
|  | smooth *Dendrodoris* | *Doriopsilla* | 39 | 0.14 | 1 |
|  | warted *Dendrodoris* | *Doriopsilla* | 84 | 0.11 | 1 |
| nad1 | smooth *Dendrodoris* | warted *Dendrodoris* | 41 | 0.33 | 0.98 |
|  | smooth *Dendrodoris* | *Doriopsilla* | 40 | 0.69 | 0.97 |
|  | warted *Dendrodoris* | *Doriopsilla* | 76 | 0.66 | 0.99 |
| nad2 | smooth *Dendrodoris* | warted *Dendrodoris* | 23 | 1.47 | 0.85 |
|  | smooth *Dendrodoris* | *Doriopsilla* | 26 | 1.96 | 0.8 |
|  | warted *Dendrodoris* | *Doriopsilla* | 56 | 2.45 | 0.86 |
| **nad3** | smooth *Dendrodoris* | warted *Dendrodoris* | 38 | 0.69 | 0.85 |
|  | smooth *Dendrodoris* | *Doriopsilla* | 36 | 0.89 | 0.73 |
|  | warted *Dendrodoris* | *Doriopsilla* | 64 | 0.85 | 0.85 |
| **nad4** | smooth *Dendrodoris* | warted *Dendrodoris* | 28 | 1.03 | 0.96 |
|  | smooth *Dendrodoris* | *Doriopsilla* | 28 | 0.99 | 0.96 |
|  | warted *Dendrodoris* | *Doriopsilla* | 69 | 0.32 | 0.98 |
| **nd4l** | smooth *Dendrodoris* | warted *Dendrodoris* | 15 | 2.03 | 0.62 |
|  | smooth *Dendrodoris* | *Doriopsilla* | 16 | 2.34 | 0.59 |
|  | warted *Dendrodoris* | *Doriopsilla* | 61 | 1.47 | 0.81 |
| **nad5** | smooth *Dendrodoris* | warted *Dendrodoris* | 32 | 1.12 | 0.91 |
|  | smooth *Dendrodoris* | *Doriopsilla* | 32 | 1.19 | 0.91 |
|  | warted *Dendrodoris* | *Doriopsilla* | 72 | 0.79 | 0.98 |
| **nad6** | smooth *Dendrodoris* | warted *Dendrodoris* | 15 | 1.31 | 0.36 |
|  | smooth *Dendrodoris* | *Doriopsilla* | 17 | 2.57 | 0.39 |
|  | warted *Dendrodoris* | *Doriopsilla* | 55 | 2.97 | 0.71 |

### Protein structure models

Sequence identity was very low when comparing smooth *Dendrodoris* with the other two Dendrodorididae morphotypes (Table 2). Nonetheless, protein models for all three main clades showed highly conserved structures for all the single genes encoded by the mitochondrial genome. However, the level of structural conservation was higher in the *COX* complex subunits than for the *NADH* or ATP synthase (*ATP*) ones. For most genes, the measure of the average distance between atoms (RMSD-values) was close to 0 Angströms (Å) (Table 2), indicating a near-perfect match between Dendrodorididae protein structures. On the other hand, the topological similarity of protein structures (TM-scores) was close to 1, supporting the similarity among Dendrodorididae structures (Table 2). The exceptions to this trend were *ATP8, ND4L,* and *NAD6* with RMSD values around 2 Å and TM-scores below 0.5 (Table 2). No apparent differences in the quaternary structure were observed when the *COX* and *NADH* protein complexes were modeled (Figure 4, Supplementary Data).

**Figure 4:**
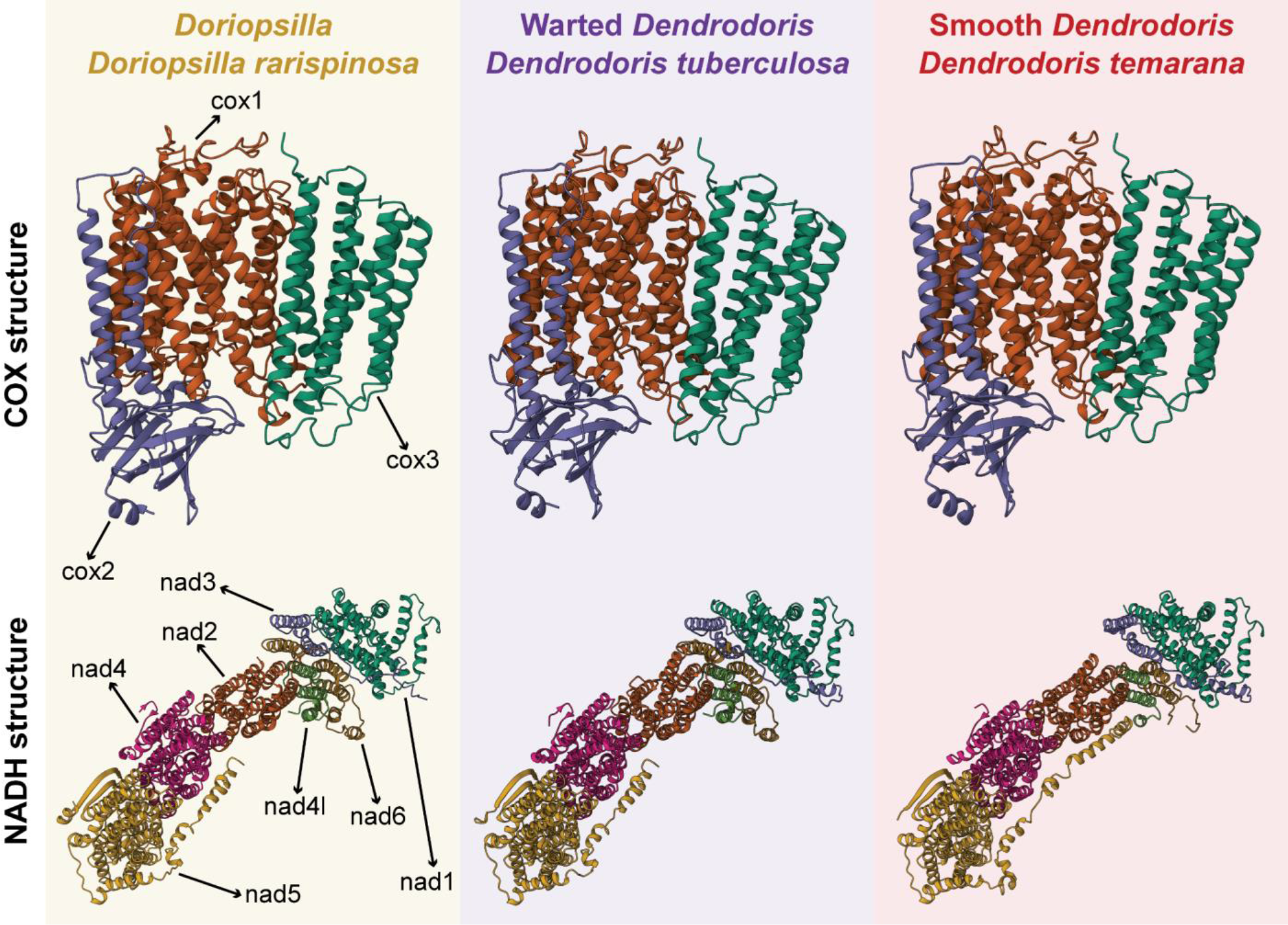
Protein models obtained for the COX and NADH complexes. Each subunit of the complex is colored in a different color.

### Nuclear signal of the family

The percentage of mapping reads was highly variable depending on the species (56.08+-0.20%, mean+- SD), with *D. tuberculosa* reaching the highest value (99.53%) and *Do. rarispinosa* the lowest (39.07%) (Table S1). After filtering, we kept a total of 30,812 SNPs for nuclear analyses. The MDS segregated each morphological variant, displaying *Doriopsilla*, the smooth *Dendrodoris,* and the warted *Dendrodoris* as independent evolutionary units. The species *De. elongata* occupied an intermediate position among the *Doriopsilla* and the other two warted *Dendrodoris* species (Figure 5A).

**Figure 5:**
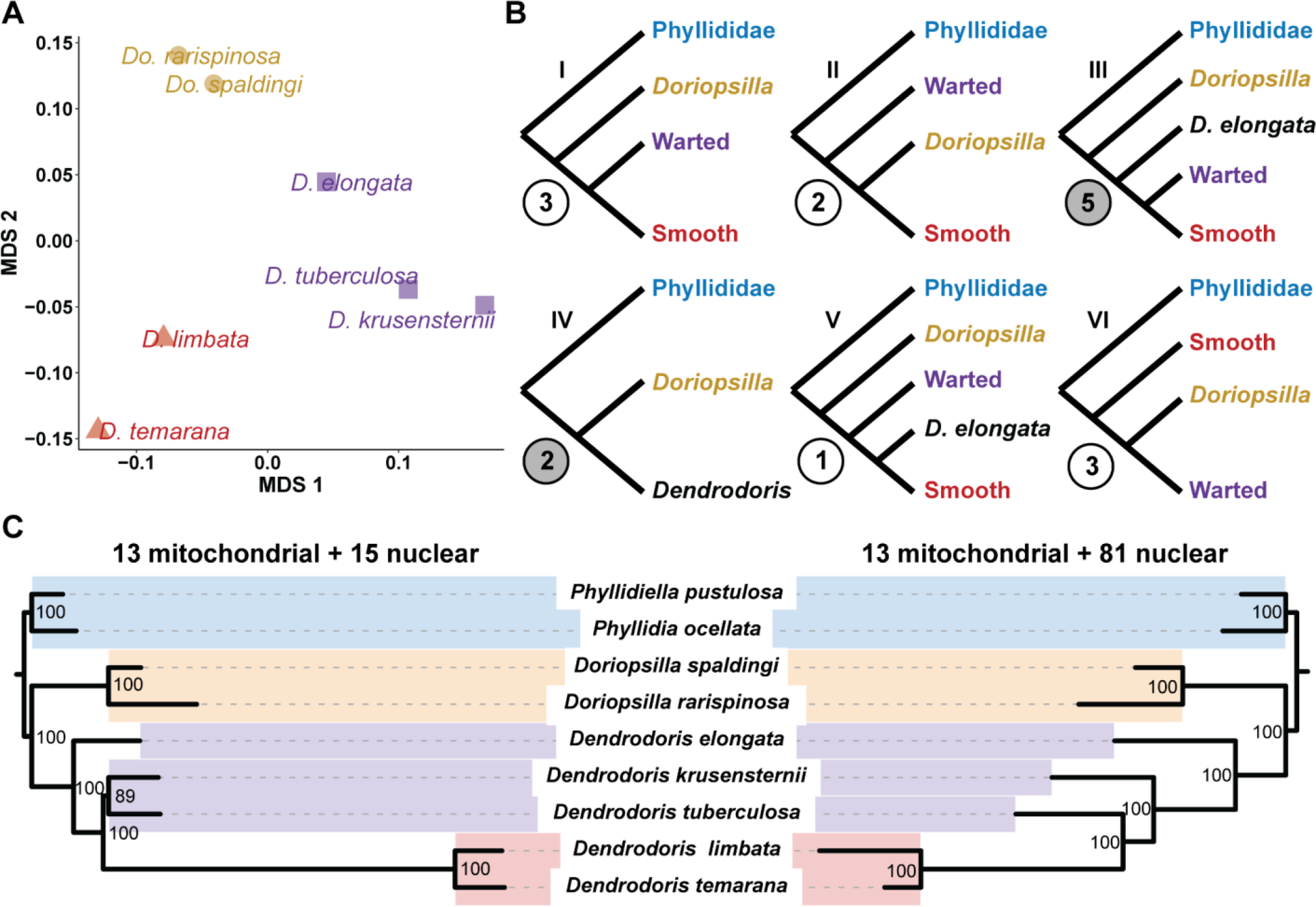
Dendrodorididae systematics. A) MDS analysis representation using nuclear information (30,812 SNPs) of the Dendrodorididae species. **B)** Different topologies were obtained with the different datasets used in the current study. Circled numbers indicate how many times each topology has been recovered. Gray circles indicate topologies retrieved when combining nuclear and mitochondrial data. **C)** Faced supermatrix topologies for the combined datasets using mitochondrial and nuclear data. Bootstrap values are reported on nodes.

### Phylogenetic analyses

#### Mitochondrial datasets

Mitochondrial trees combining CDS and rRNA data and including different codon positions, presented topological differences depending on the dataset used for the analysis (Table S2, Figure 5b). However, all of them displayed a long branch for all smooth *Dendrodoris* species (Figure S1), with low bootstrap node values (BS) of BS=48 when using all three positions of each codon, BS=23 for the first and second positions, BS=50 when using only the second position, and BS=36 when only using the first position. When using all positions of the codon, *Doriopsilla* was the sister group to the smooth *Dendrodoris* clade, and warted *Dendrodoris* were the sister group of the remaining members of the family Dendrodorididae. Using the first and second positions of the codon, or only the second position, the clade of warted *Dendrodoris* was sister to *Doriopsilla*, and the smooth *Dendrodoris* clade formed a basal polytomy with the outgroup. Finally, using only the first position of the codon, the genus *Dendrodoris* was monophyletic, with a long branch separating smooth and warted *Dendrodoris*, and sister to *Doriopsilla*.

#### Nuclear datasets

The nuclear analyses produced two topologies (Table S2, Figure S2). All topologies supported monophyly of smooth *Dendrodoris* and paraphyly of warted *Dendrodoris*, with *D. elongata* as the sister clade to the remaining *Dendrodoris*. They disagree in considering *D. tuberculosa* and *D. krusensternii*, either as a grade (81 BUSCO Supermatrix) or as a clade (remaining analyses), sister to smooth *Dendrodoris*. *Doriopsilla* was supported as a sister taxon to all *Dendrodoris* in both topologies. All supermatrix nodes had bootstraps (BS) above 90, except for the 15 gene dataset where the support for smooth *Dendrodoris* monophyly was BS=51 and for warted *Dendrodoris* BS=55. The 81 nuclear genes ASTRAL supertree recovered full support (1) for all nodes. Conversely, the 15 nuclear genes ASTRAL tree had more variable, albeit high, node support values, namely 0.87 for smooth *Dendrodoris*, 0.93 for warted *Dendrodoris*, and 1 for the remaining nodes.

#### Combined datasets

The combined datasets inferred highly supported phylogenies, although different topologies were found (Table S2, Figure 5c). Supermatrix node support was higher than supertree node support (Figure S3), with bootstrap values of 100 in all nodes. The only exception was the concatenated dataset including the 13 mitochondrial + 15 nuclear genes, which provided a bootstrap value of 89 for thewarted Dendrodoris clade.

## DISCUSSION

The results presented here are novel in the study of structural rearrangements and accelerated mutation events in nudibranchs. The rearrangement of the mitogenome of smooth *Dendrodoris* species and the elevated mutation rates produced molecular changes in the genes contributing to the respiratory machinery of this group. However, we have corroborated that protein structure is conserved overall, and therefore, functionality is expected to be maintained. Nevertheless, the high mutation rates in mitochondrial genes had a relevant effect on the group’s phylogeny, and is the major responsible for the previously observed mito-nuclear discordance, resolved in the present study.

### Mitochondrial gene reorganization and mutation rates

Structural rearrangements are hypothesized to result from duplication events followed by the random deletion of one of the copies (Shi *et al*, 2015; Françoso *et al*, 2023; Ghiselli *et al*, 2021). Although mollusks are well known for having large and structurally diverse mitochondrial genomes (Ghiselli *et al*, 2021), dorid nudibranchs are supposed to be an exceptional case, as they are characterized by short compact mitogenomes (Do *et al*, 2022). This mitogenomic structure is thought to be relatively stable (Grande *et al*, 2008; Xie *et al*, 2019) due to a high degree of gene overlap that could accentuate the effect of deleterious mutations and/or rearrangements (Ghiselli *et al*, 2021). Thus, as mutations may simultaneously affect multiple genes, natural selection would quickly remove them, maintaining the original mitogenomic structure. Although mitogenome arrangements have been already reported in nudibranchs, specifically in the genus *Hypselodoris (Do et al, 2022)*, the extremely fast mutation rates suggested by the extremely high dN and dN/dS values in smooth *Dendrodoris* challenge traditional views on mitogenome evolution (Karagozlu *et al*, 2016; Varney *et al*, 2021). It is worth mentioning that the increase in mutation rate is likely to have occured after the split of the family Dendrodorididae from its sister group Chromodorididae, which is estimated to have happened 40 million years ago (https://timetree.org/). Given this time window and the high d*N* and d*N*/d*S* values found for congeneric *Dendrodoris* species, their mitochondrial genome mutation rates are the most extreme cases found to date in mollusks. Our average d*N*/d*S* values of 0.273 surpass by far the highest molluscan rate reported so far of d*N*/d*S*=0.161, corresponding to the asexual snail *Potamopyrgus antipodarum* (Neiman *et al*, 2010). Smooth *Dendrodoris* presented significantly higher d*N*/d*S* values than the other Dendrodorididae species in most of the genes. Most of the d*N*/d*S* values were below 1, thus indicating purifying selection (Wang *et al*, 2021). The exceptions to this trend were comparisons in the genes *atp8* (*D. temarana* vs. *P. pustulosa;* d*N*/d*S*=1.19), *nad4l* (*D. temarana* vs. *D. elongata*; d*N*/d*S*=4.12), and *nad6* (*D. temarana* vs. *P. ocellata*; d*N*/d*S*=34.63, and *D. limbata* vs. *D. elongata* d*N*/d*S*=1.28 ), which might indicate potential positive selection. Nevertheless, the high d*S* values found on most of the genes of smooth *Dendrodoris* suggests that *Dendrodoris* sequences might be highly saturated (Shapiro & Alm, 2009), preventing the detection of positive selection through d*N*/d*S* ratios on the current mitochondrial gene sequences. Therefore, a positive selection event occurring in the past should not be ruled out (Shapiro & Alm, 2009).

### Protein models validate orthology

The unprecedented prevalence of non-synonymous mutations between sibling taxa in the family Dendrodorididae and the high number of indels in all 13 CDSs + 2 rRNAs is unique among heterobranch sea slugs. Although some authors have suggested on *cox1* and *16S* this could be the result of the sequencing of pseudogenes or exogenic contamination (Hallas *et al*, 2017; Korshunova *et al*, 2020), our protein subunits and protein complexes models confirm that despite having non-synonymous mutations, the protein structure is maintained. Therefore, *Dendrodoris cox1,* and by extension also 16S genes, most likely do not represent deleterious pseudogenes but functional orthologous genes. The only proteins not modeled correctly are the smallest subunits of each complex, whose deleterious effect on the entire protein complex might be negligible. Because COX and NAD protein subunits interact with each other, mutations in either subunit can lead to different evolutionary scenarios. On the one hand, one of the interacting subunits exerts selective pressure on the newly generated mutation in the second subunit, and it is purified. On the other hand, folding proteins such as chaperones reduce the harmful effects of the subunit mutation, and the accumulation of compensatory mutations can occur. (Hill, 2020; Maisnier- Patin *et al*, 2005). This effect can lead to a cascade of mutations in all proteins involved in the complex, sweeping surrounding regions (whole mitogenomes in the present study), accelerating the mutation and selection processes (Hansen, 2013; Paixão & Barton, 2016) through synergistic epistasis. As a result, extremely high evolutionary rates can occur, such as reported in marine copepods (Stern *et al*, 2022). However, the mutagenesis of this effect in mollusks is still poorly understood and neglected in nudibranchs. Future studies using reference genomes of Dendrodorididae species with a broader taxon sampling should shed light on these processes.

### Dendrodorididae systematics and evolution: mitogenomes

Our protein models confirm that phylogenetic studies can be safely carried out when using mitochondrial markers, as these genes are most likely orthologous. Our phylogenetic results using mitochondrial data validate the presence of a long branch separating the smooth *Dendrodoris* clade (Hallas *et al*, 2017; Korshunova *et al*, 2020). In all cases, it has a low node support value, suggesting that the long branch reflects high genetic distances between taxa with unreliable phylogenetic relationships (Bergsten, 2005; Kück *et al*, 2012). Although we cannot establish a reliable phylogeny based on mitochondrial data due to topology inconsistencies, sufficient indicators support the hypothesis that smooth and warted *Dendrodoris* are potentially two reciprocal monophyletic evolutionary units. Among evidence, we found that mitochondrial genetic distance is exceptionally high, and the presence of highly divergent mitogenome structures involving indels between groups which can be considered as an autapomorphic syntenic trait (Schreeg *et al*, 2016). The difference in the mitogenome size of ∼1000 bp and a completely different GC content: x̄ =28.10% in smooth and x̄ =35.23% in warted *Dendrodoris*, is consistent with morphological differences, and further corroborates that these two groups of *Dendrodoris* should be regarded as different genera. AT-rich mitogenomes have been described for deep-sea invertebrates compared to their shallow-water relatives (Cejp *et al*, 2022; Yang *et al*, 2019). Because three deep-sea *Dendrodoris* species inhabit habitats below 100 meters depth (Valdés, 2001), in contrast to the shallow depths of most *Dendrodoris* (10–20 meters for warted and ∼0 meters for smooth, respectively), the inclusion of these deep-sea species in future studies could provide a link between smooth and warted clades that is crucial for clarifying the evolutionary story of the family Dendrodorididae.

The mitogenome rearrangement and high mutation rates on the smooth *Dendrodoris* lineage represents a case of extremely fast evolution. In other groups of organisms (e.g., sauropsid reptiles), a direct correlation between rapid speciation events and accelerated evolutionary rates was found in mitochondrial genomes (Eo & DeWoody, 2010). This situation could be reflected in the biodiversity of each *Dendrodoris* morphotype. While warted *Dendrodoris* only has about ten species, the smooth *Dendrodoris* lineage accounts for approximately 30 species. However, both lineages might be more diverse since *Dendrodoris* has been found to contain several cryptic species (Galià-Camps *et al*, 2022). Studies conducted in orthopteran insects suggest that mitogenome evolutionary rates are constrained by energy-related selection (Chang *et al*, 2020) and concluded that grasshopper species subjected to weaker selection pressures on respiration had faster substitution rates than those subjected to highly demanding respiration processes. Warted *Dendrodoris* are commonly found below 20 meters. In contrast, smooth *Dendrodoris* are found under rocks in shallow waters between 0 and 5 metres depth where oxygen levels are higher, which is consistent with the respiratory constraint release hypothesis. In this context, the ecological niche that occupies smooth *Dendrodoris* characterized by high oxygen levels, food resources, and microhabitats, could have promoted speciation through niche specialization, resulting in the adaptive radiation of shallow water smooth *Dendrodoris* species.

### Nuclear data confirms mitogenome accelerated mutation rates

The generation of the first genome draft of seven Dendodorididae species has been a keystone to unveil the nuclear processes underlying Dendrodorididae evolution. It is noteworthy that Dendrodorididae genomic resources have to be improved, since *D. tuberculosa*, our best pseudo-reference assembly, only had 51% of BUSCO genes, thus indicating that approximately half of the genome information is missing. In front of the enlightening results obtained with partial genomic data, the potential of chromosome-level assemblies in evolutionary molluscan studies is remarkable. The inclusion of genome-wide SNPs and sequences from nuclear coding genes has been crucial for validating the dichotomy between smooth and warted *Dendrodoris* species. The SNP data support the differentiation between smooth and warted *Dendrodoris*, as they split on the first axis (Figure 4a), and both groups are equally distinct from the closely related genus *Doriopsilla*. Similarly, the phylogenetic position of smooth *Dendrodoris* is firmly supported when using nuclear data or when combining it with mitochondrial data since low mutation rates of nuclear markers allow better inferences of deeper phylogeny nodes (Talavera & Vila, 2011). However, including more nuclear genes in the analysis directly impact the topology and, thus, the clusters generated. Our results support two alternative topologies. The first topology, supported by 81 BUSCO + 13 mitochondrial genes, supports the monophyly of the genus *Dendrodoris*. The second one, supported by SNP data and the 15 BUSCO + 13 mitochondrial genes, indicates that *Dendrodoris* could have split into three lineages. The limited taxon sampling prevented us from choosing among alternative topologies. Nonetheless, the latter hypothesis conciliates with morphological characters, as it splits smooth and warted *Dendrodoris,* and places *D. elongata* as sister to *Doriopsilla*. *Dendrodoris elongata* has small tubercles, typical from warted *Dendrodoris*, but presents a softly spiculated mantle in line with *Doriopsilla* species (Furfaro *et al*, 2022).

We encourage future studies to expand the taxon sampling with additional individuals for each morphotype and ecotype to fully resolve the systematics of the family Dendrodorididae, and reveal the selective pressures that led the family to rumble the classical interpretation of the neutral mutation theory and its evolution.

## CONCLUSIONS

Our study reveals accelerated evolutionary changes in the mitogenome of the nudibranch genus *Dendrodoris*, the second case documented in a nudibranch. Although the mechanisms behind this process and their implications for respiration remain unknown, the habitat shifts might be a potential trigger for this process. Future studies should include reference genome assemblies for multiple species of Dendrodorididae occupying different ecotypes to identify the associated mitochondrial and nuclear genes. Additionally, reference genomes could fully resolve the phylogenetic relationships within the family. Our study represents a first attempt to explore the history of the family Dendodorididae through their mitochondrial evolution, a neglected area in malacology but with far-reaching implications for many other taxa with diverse habitats and ecologies.

## MATERIAL & METHODS

### Taxon sampling

Seven Dendrodorididae species were collected in shallow waters of NE Spain (Catalonia), USA (California and Hawaii) (Table S3). The sampling was designed to obtain at least two representatives of each of the main morphotypes in the family: smooth *Dendrodoris* (*D. limbata* and *D. temarana*), tuberculated *Dendrodoris* (*D. tuberculosa*, *D. elongata* and *D. krusensternii*) and *Doriopsilla* (*D. rarispinosa* and *D. spaldingi*) (Figure 1).

### DNA extraction and library preparation

DNA was extracted from a ∼25mm^2^ fragment of the foot of a specimen of *Dendrodoris temarana* using a customized phenol-chloroform protocol using magnetic beads based on (Mayjonade *et al*, 2016). The DNA extraction was sent to Novogene (UK) for short-read Illumina genome sequencing. The genomic library was prepared with an insert size of 350 bp, and an Illumina NovaSeq 6000 platform (San Diego, CA) was used to sequence 150 bp paired-end reads, aiming for 60 Gb output. DNA of the remaining specimens was extracted from foot fragments of ∼25mm^2^ using the Gentra Puregene Tissue Kit (QIAGEN) by following the manufacturer’s instructions. An initial amount of DNA ranging from 300 to 500 ng was used for library construction using all standard kit reagents from the Illumina DNA prep (M) Tagmentation (Illumina, San Diego, CA), altogether with Illumina commercial indexes (IDT for Illumina DNA/RNA UD Indexes set A, Tagmentation) by following the kit manual instructions. Library preparations (insert size: 350 bp) were sent to Macrogen (Macrogen Korea) for 150 bp paired-end reads sequencing on a HiSeqX platform with 14 Gb output each.

### Data curation, assembly, and annotation

Raw data was submitted to the European Nucleotide Archive (ENA) under the Bioproject PRJEB75450. We discarded low-quality data and adapters using Trimmomatic 0.39 (Bolger *et al*, 2014), applying default parameters. For the mitochondrial genome assembly reconstruction, NOVOPlasty 4.3.1 (Dierckxsens *et al*, 2017) was run for all the individuals using a k-mer size of 21 and the *cox1* gene sequence of each species as seed (Table S3), except for *De. elongata* as it was unavailable in public databases and we used the *D. krusensternii cox1* instead. The complete mitogenome of *Phyllidia ocellata (Xiang et al, 2016)* (NCBI Accession number = KU351090) was used as the reference mitogenome for all assemblies (Table S3). The software MITOS2 (Donath *et al*, 2019), implemented in GALAXY web https://usegalaxy.eu/, was used to annotate ribosomal RNAs (rRNA), coding sequences (CDS), and transfer RNAs (tRNA), which were subsequently visualized and manually curated using Geneious Prime 2022.1.1 (Biomatters Ltd). SPAdes-3.11.0 (Prjibelski *et al*, 2020), was run on the filtered reads with default parameters to obtain a *de novo* assembly of the nuclear genome for each species. All nuclear and mitochondrial assemblies were submitted to ENA as part of the Bioproject PRJEB75450.

### Mitogenome and gene characteristics

For comparison, we retrieved two additional mitogenomes from Genbank belonging to the family Phyllididae: *Phyllidia ocellata* and *Phyllidiella pustulosa* (Table S3, Figure 1). Mitogenome length and GC content were calculated with Geneious Prime. Mitogenomic sequences were aligned with MAFFT 7.376 using the L-INS-i method (Katoh & Standley, 2013) and visualized and curated in Geneious Prime. Subsequently, we compared the gene order across the mitogenome alignment. Additionally, each individual rRNA (as nucleotide) and CDS (as amino acid) was aligned separately using MAFFT with a posterior manual curation. CDS nucleotide alignment was based on codon position and carried out with the software PAL2NAL v14 (Suyama *et al*, 2006), using as reference CDS amino acid alignments. We conducted a pairwise comparison among aligned nucleotide CDS sequences to evaluate the number of non-synonymous mutations per non-synonymous site (d*N*), synonymous mutations per synonymous site (d*S*), and their ratio (d*N*/d*S*), with the Maximum Likelihood method implemented in the codeml function of PAML 4.10.1 (Yang, 2007). For each comparison, we used a free-ratio “branch” model which allows the calculation of a distinct ratio for each branch, equilibrium codon frequencies as free parameters (codonfreq = 3), and only considered those positions without gaps to avoid ambiguous data (cleandata = 1). General Linear Models (GLM) to evaluate the factors influencing d*N*, d*S*, and d*N*/d*S* were implemented in R considering: gene (*atpX*, *coxX*, *nadX*), sequence pairwise comparison [involving or not involving smooth *Dendrodoris* (see results)], and their interaction. Both d*N* and d*N*/d*S* values were log10 transformed. To further evaluate factors found to be significant, we conducted a pairwise post hoc analysis with a Tukey correction using the function “contrast” of the R package emmeans (Lenth *et al*, 2020). We exported the results as graphs using the R package ggplot2 (Wickham *et al*, 2016).

### Mitogenome protein structure models

To evaluate the functionality of the annotated CDS, we modeled and compared the protein structure of each CDS among Dendrodorididae morphotypes. Based on the GMQE score, we built homology models with SWISS-MODEL (Waterhouse *et al*, 2018) using the most similar available protein to our amino acid sequence as a template. We repeated the same procedure for cytochrome oxidase (*COX*) and nicotinamide adenine dinucleotide (*NAD*) protein complexes to validate no aberrant structure in the protein complex assembly. This approach was not carried out for the ATP synthase (ATP) protein complex since only two small subunits are synthesized in the mitochondria and therefore, the protein complex model remained largely incomplete. The resulting models were pairwise-compared between species, using the pairwise structure alignment tool with the algorithm jFATCAT (rigid) of RCSB PDB (https://www.rcsb.org/) (Duarte *et al*, 2022) to test the percentage of sequence similarity, the Root-mean- square deviation of atomic positions (RMSD-value), and the template modeling (TM)-score (Zhang & Skolnick, 2004; Kufareva & Abagyan, 2011).

### Mitochondrial phylogenies

To generate the mitochondrial phylogenies, we followed both a supermatrix and a supertree approach for both amino acid and nucleotide alignments. The supermatrix contained all CDS and the two rRNA for the 7 Dendrodorididae and the two Phyllididae species. Maximum likelihood analyses were performed with IQ-TREE2 (Minh *et al*, 2020). No partition was defined when conducting amino acids. For nucleotide sequences, we ran the following approaches: 1) without partitions, 2) we partitioned each codon position per CDS + rRNA, 3) we partitioned the first and second position of the codon per CDS + rRNA, 4) we only used the first position of the codon per CDS + rRNA, and 5) we only used the second position of the codon per CDS + rRNA (Table S2). For the supertree, we performed maximum likelihood analyses for each CDS and rRNA without defining partitions regardless of amino acid or nucleotide sequence, and we obtained a consensus tree using ASTRAL 5.7.7 (Table S2) (Zhang *et al*, 2018). Phylogenies were visualized and customized in iTOL 5 web server (Letunic & Bork, 2021).

### Nuclear and combined phylogenies

Completeness regarding conserved metazoan single-copy orthologs (metazoa_odb10) was assessed with BUSCO 5 (Manni *et al*, 2021). Single-copy ortholog gene coordinates were provided by BUSCO, and were posteriorly used to retrieve the sequences from the genome assemblies of each species with gffREAD v.0.12.8 (Pertea & Pertea, 2020), and ortholog relationships were assessed using OrthoFinder 2.5.4 using default parameters. Ortholog sequences were aligned and manually curated to avoid paralogs and non-reliable sequences. Finally, we kept gene alignments with representatives from all species (complete dataset) and those including at least four different species (partial dataset). For nuclear phylogeny, we analysed the supermatrix with IQTREE 2 and a supertree with ASTRAL for both complete and partial species datasets, without partitions (Table S2). We then combined the mitochondrial dataset with the respective nuclear datasets to obtain a dataset without missing data comprising 13 mitochondrial and 15 nuclear genes, and one with missing data (5 out of 7 species had the sequence) comprising 13 mitochondrial and 81 nuclear genes (Table S2). Supermatrix and supertrees were obtained as previously reported. The resulting phylogenies were visualized and customized with iTOL 5 web version.

### Nuclear genetic distance signaling

We evaluated which genome assembly was best suited to be used as a pseudo-reference genome to generate a catalog of genetic variants among species. Our evaluation was based on which genome assembly better resolved the trade-off between total length and BUSCO completeness to be used as a pseudo-reference genome for genotyping. We mapped the clean reads of the seven Dendrodorididae species to the pseudo-reference (*D. tuberculosa*, see results) with bwa 0.7.1 mem (Li & Durbin, 2009), and genotypes were called using the command mpileup in bcftools 1.16 (Danecek *et al*, 2021). Vcftools 0.1.17 (Danecek *et al*, 2011), was used to keep biallelic SNPs present in at least 5 of the 7 species, with a minimum locus depth of 5 reads, a maximum locus depth of 100, and a minimum allele frequency of 15% to discard species-specific loci. PLINK 1.9 (Purcell et al. 2007) was used to perform a multidimensional scaling (MDS) on the data, and the results were plotted with ggplot2 (Wickham, 2009) in R.

## DATA AVAILABILITY

Raw data and genome assemblies have been uploaded to the European Nucleotide Archive (ENA) under the BioProject PRJEB75450. Structural models, alignments, genotyping datasets, and mitogenome annotations have been provided in a repository hosted in github (https://github.com/CGaliaCamps/Dendrodorididae_mitogenomes).

## AUTHOR CONTRIBUTIONS

**Carles Galià-Camps:** Conceptualization, Data Curation, Methodology, Software, Validation, Formal analysis, Investigation, Writing – original draft preparation, Writing – review and editing, Visualization, Project administration. **Tilman Schell:** Software, Data Curation, Validation, Resources, Writing – review and editing, Supervision. **Alba Enguídanos:** Methodology, Validation, Investigation, Resources, Writing – review and editing, Supervision. **Cinta Pegueroles:** Validation, Resources, Writing – review and editing, Supervision. **Miquel Arnedo:** Validation, Resources, Writing – review and editing, Supervision, Funding acquisition. **Manuel Ballesteros:** Conceptualization, Resources, Writing – review and editing, Supervision, Funding acquisition. **Ángel Valdés:** Conceptualization, Methodology, Validation, Resources, Writing – review and editing, Supervision, Funding acquisition. **Carola Greve:** Conceptualization, Methodology, Validation, Investigation, Resources, Writing – review and editing, Supervision, Funding acquisition

## ACKNOWLEDGEMENTS

This project was funded by the Centre for Translational Biodiversity Genomics (LOEWE-TBG) through the program LOEWE–Landes-Offensive zur Entwicklung Wissenschaftlich-ökonomischer Exzellenz of Hesse’s Ministry of Higher Education, Research, and the Arts (HMWK). Additional funding was provided by grant [PID2019-105794GB] from Ministerio de Economía y Competitividad of Spain, and the grant [PID2020-118550RB] from the Ministerio de Ciencia e innovación of spain. Partial funding was also granted with the U.S. National Science Foundation [DEB-1355177]. MA and AE are part of the [2021- SGR689], and CGC and MB are part of the [2021 SGR 01271], both funded by the Agència de Gestió d’Ajudes Universitàries of the Catalan government.

## SUPP TABLES CAPTIONS

**Table S1: Sequencing, Mitochondrial assembly, Nuclear assembly, genes used for phylogeny, and SNP data features.** For each species, we provide raw reads (Gb), polished reads (Gb), mitochondrial assembly size (pb), mitochondrial coverage, nuclear assembly size, complete, fragmented and missing BUSCO genes, number of mitochondrial markers used in phylogenies, BUSCO used allowing missing data, and percentage of reads mapped to the nuclear assembly of *Dendrodoris tuberculosa*

**Table S2: Phylogenetic approaches summary.** For each approach we provide the name of the model, number of genes included, genomic compartment of the genes, sequence type, analysis performed, parition used and topology obtainded (see Fig. 4b)

**Table S3: Samples used for the study.** For each species we provide latitude and longitude of collection site, and reference *cox1* accession number

**Supplementary Spreadsheet 1: Tuckey post-hoc conducted on GLM significant interactions.** For each pair- wise comparison, we provide Gene, Comparison (involving or not smooth *Dendrodoris*), Comparison being tested (Test), Estimate, Standard Error, Degrees of Freedom (DF), t-ratio, and p-value. Nothe that asterisks indicate that this factor is being pairwised tested. Significant p-values are hihglighted in bold.

**Figure S1:**
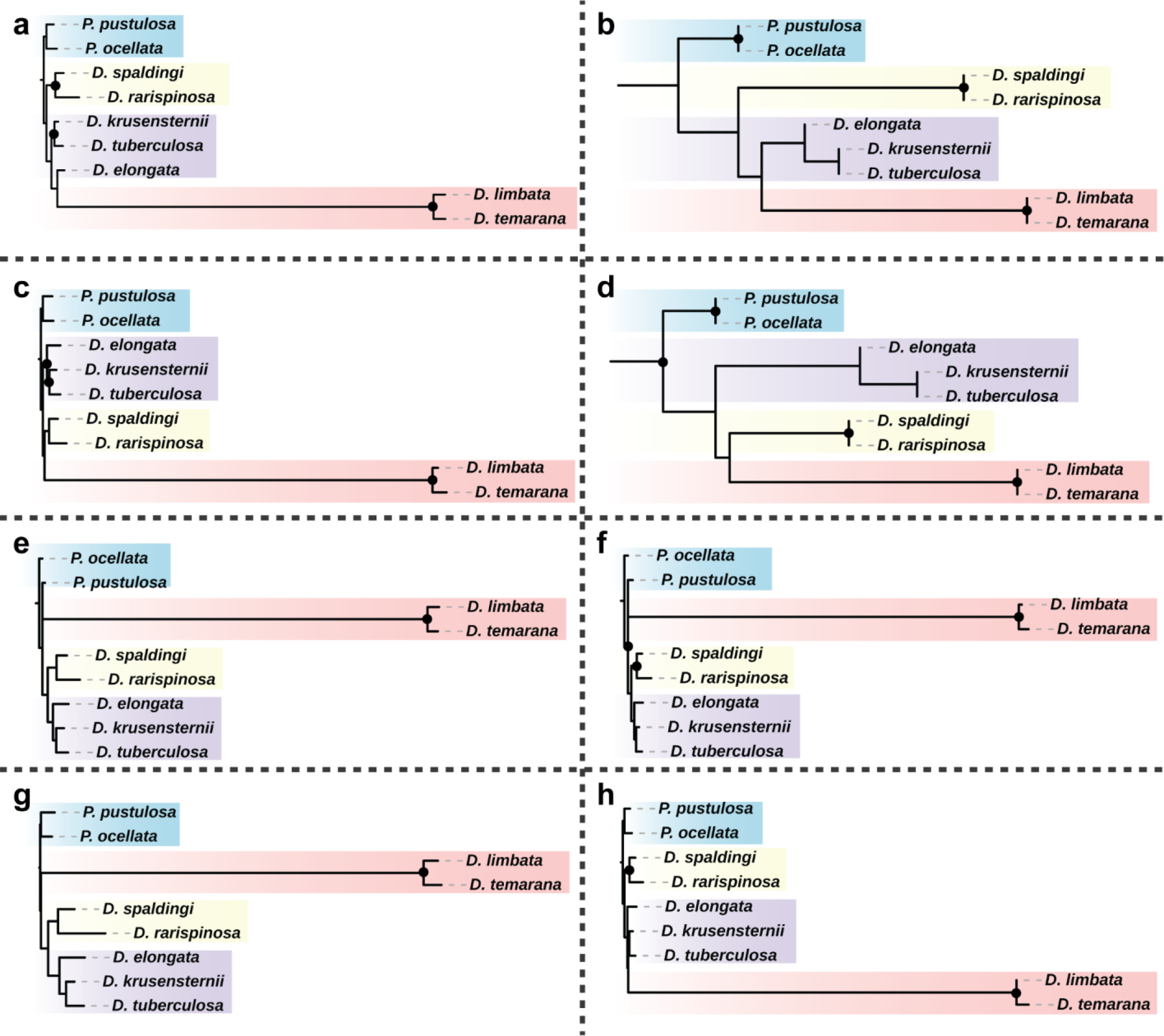
Mitochondrial phylogenies obtained using different datasets. Nodes with Bootstrap (BS) and ASTRAL node support >95 are indicated with a dot. **a)** Supermatrix maximum likelihood (ML) approach using amino acid sequences of all CDS. **b)** Supertree approach using amino acid sequences of all CDS. **c)** Supermatrix ML approach using nucleotide sequences from all CDS + rRNA without partitions. **d)** Supertree approach using nucleotide sequences from all CDS + rRNA without partitions. **e)** Supermatrix ML approach using nucleotide sequences from all CDS + rRNA partitioning each position of the codons. **f)** Supermatrix ML approach using nucleotide sequences from all CDS + rRNA partitioning the first and second positions of the codons. **g)** Supermatrix ML approach using nucleotide sequences from all CDS + rRNA partitioning only the second positions of the codons. **h)** Supermatrix ML approach using nucleotide sequences from all CDS + rRNA partitioning only the first positions of the codons.

**Figure S2:**
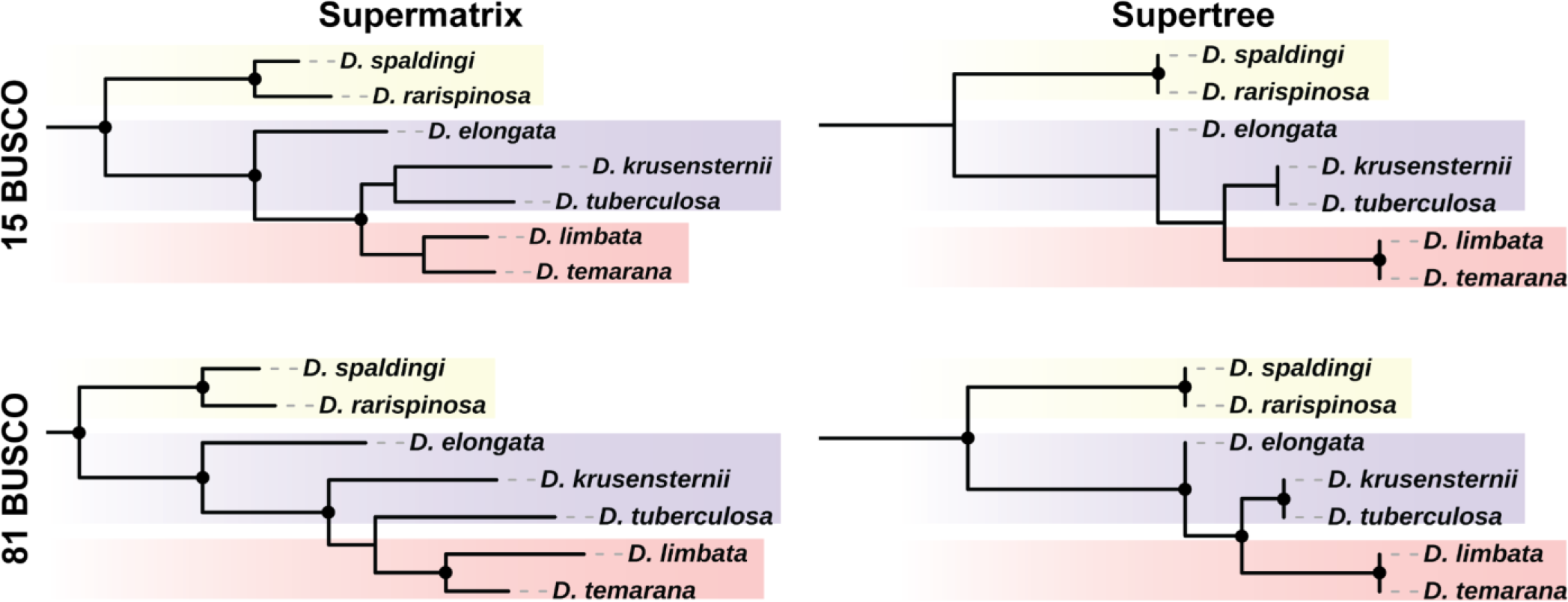
Nuclear phylogenies obtained using different datasets. Top-row phylogenies use a dataset including 15 BUSCO genes without missing data. Bottom-row phylogenies use a dataset including 81 BUSCO genes present in at least 5 species. Left-column phylogenies use a supermatrix ML approach. Right-column phylogenies use a supertree approach. All approaches are based on amino acid sequences. Nodes with Bootstrap (BS) and ASTRAL node support >95 are indicated with a dot.

**Figure S3:**
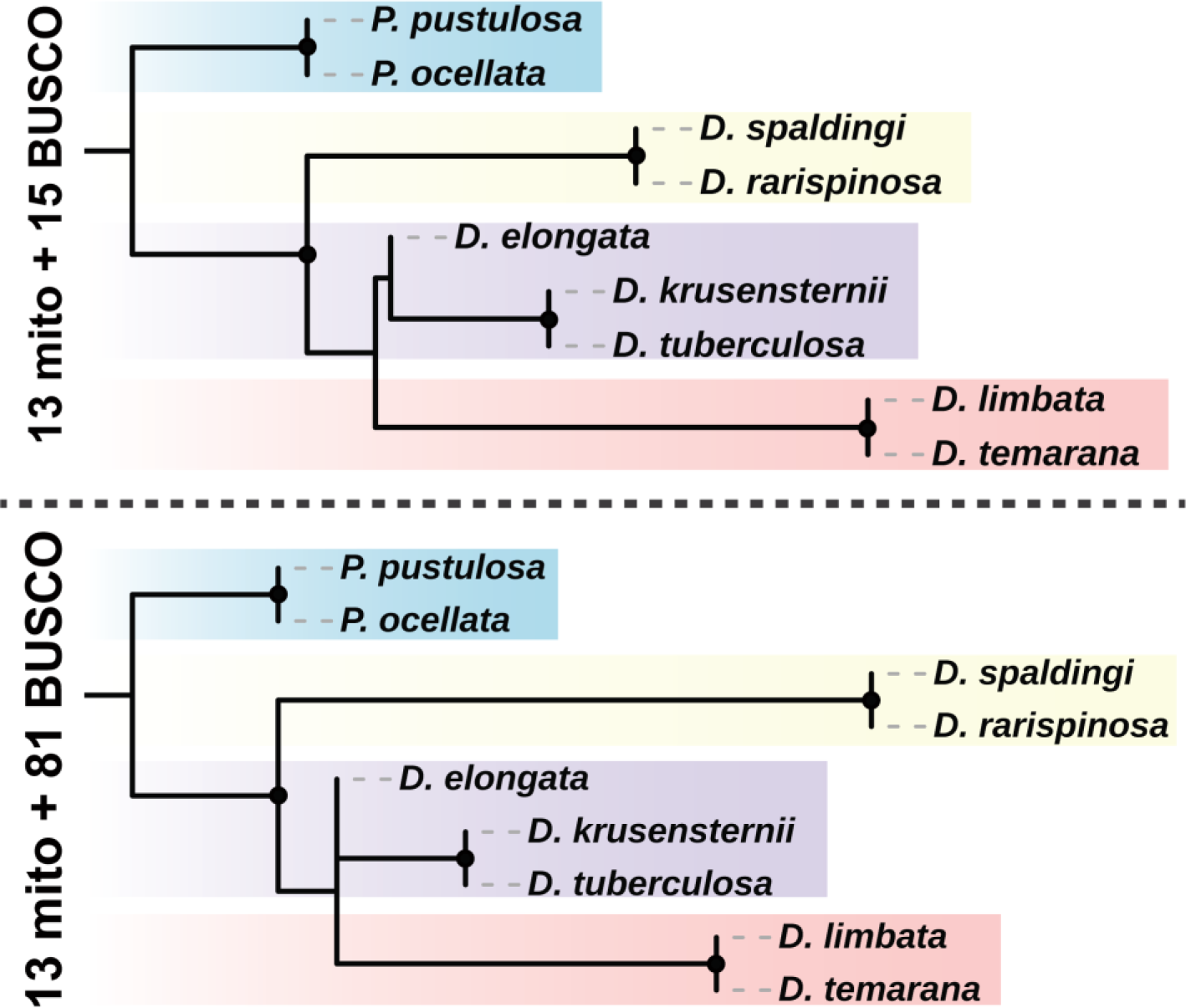
Concatenated phylogenies obtained using different datasets. The top phylogeny is based on a supertree approach using amino acid sequences of all mitochondrial CDS and 15 BUSCO genes without missing data. The bottom phylogeny is based on a supertree approach using amino acid sequences of all mitochondrial CDS and 81 BUSCO genes present in at least 5 Dendrodorididae species. All approaches are based on amino acid sequences. Nodes with Bootstrap (BS) and ASTRAL node support >95 are indicated with a dot.

